# Methods for Evaluating Effects of Transgenes for Quantitative Traits

**DOI:** 10.1101/2022.10.22.513367

**Authors:** Julien F. Linares, Nathan D. Coles, Hua Mo, Jeffrey E. Habben, Sabrina Humbert, Carlos Messina, Tom Tang, Mark Cooper, Carla Gho, Ricardo Carrasco, Javier Carter, Jillian Wicher Flounders, E. Charles Brummer

## Abstract

Transgenes that improve quantitative traits have traditionally been evaluated in one or a few genetic backgrounds across multiple environments. However, testing across multiple genetic backgrounds can be equally important to accurately quantify the value of a transgene for breeding objectives. Creating near-isogenic lines across a wide germplasm space is costly and time consuming, which renders it impractical during early stages of testing. In this experiment, we evaluate three approaches to sample the genetic space while concurrently testing across environments. We created both transgenic and non-transgenic doubled haploid lines, F_2:3_ lines, and bulk F_3_ families to determine if all methods resulted in similar estimation of transgene value and to identify the number of yield trial plots from each method necessary to obtain a stable estimate of the transgene value. With one exception, the three methods consistently estimated a similar effect of the transgene. We suggest that bulked F_3_ lines topcrossed to a tester inbred is the most effective method to estimate the value of a transgene across both genetic space and environments.

## INTRODUCTION

Quantitatively inherited traits are controlled by multiple quantitative trait loci (e.g., Lynch and Walsh, 1998; Falconer and Mackay, 1996). Generally, quantitative trait loci (QTL) for complex traits such as grain yield of crop plants can have drastically different effects depending on the genetic background evaluated (Kramer, 2009; Cheng et al., 2012; Powell et al., 2021). For instance, in maize, the detection and effect size estimates of QTL identified using two different tester inbreds showed considerable differences for grain yield but not for other traits such as plant height and grain moisture (Melchinger et al., 1998). In part because of the lack of a predictable response due to significant QTL × genetic background interactions, large effect QTLs for complex traits are frequently published but have rarely been effectively deployed in breeding programs to develop commercialized cultivars (Bernardo, 2008).

For transgenes intended to target complex traits, the limited research to date points to similar complexities (e.g,. Simmons et al., 2021). Transgenes targeted to affect quantitative traits have characteristics in common with large effect QTL, where the observed effect depends on the genetic backgrounds in which it is evaluated. In the maize commercial seed industry, doubled haploids (DH) have become the primary method of line development over the past 20 years (Chaikam et al., 2019). Transgene evaluation for commercial assessment is typically done by comparing one or a few near-isogenic lines that may not provide a clear picture of the effect of a transgene targeting a quantitative trait across the breadth of a germplasm under improvement in a breeding program. Improving the accuracy of estimating the value of a transgene for quantitative traits across the breeding germplasm would require an expansion of transgene evaluation beyond simple near-isogenic line testing.

To expand testing, breeders could randomly sample germplasm from throughout the breeding pool, develop numerous DH inbred lines with and without the transgene, and then use the contrasting groups of lines to determine the transgene value and stability of expression for the target quantitative traits across germplasm backgrounds. While this traditional transgene testing approach would provide the desired information on transgene value, expanding this testing program across the breeding germplasm would require a significant investment of time and resources. Therefore, it is impractical to upscale early in the testing pipeline, especially when many potential transgenes targeting multiple quantitative traits are being considered for commercialization.

Alternative testing methods could possibly enable an assessment of the breeding potential of a quantitative trait transgene throughout the program’s germplasm while also optimizing time and testing resources. In this experiment, we compared the use of doubled haploid lines for assessing transgene value with F_2:3_ lines and F_3_ families. Although the DH pipeline is the quickest approach for inbred development, creating doubled haploids requires more time than simple F_2:3_ or F_3_ line generation methods, is resource intensive, and requires many test plots to evaluate both transgenic and non-transgenic individual lines across multiple environments. Although F_2:3_ lines are less widely used in commercial maize breeding programs, and they are not fully homozygous, they enable testing of population level transgene estimates within two inbreeding generations. Significant testing resources are still required to evaluate sets of transgenic and non-transgenic F_2:3_ lines per family. Finally, we evaluate a third method of testing F_3_ bulks with and without the transgene. For F_3_ bulks, the within population variance would be captured in a single plot and estimates of the transgene effect could be done across two plots, one having the quantitative trait transgene of interest and the other as the null comparator, facilitating rapid evaluation of the average effect of the transgene across many different genetic backgrounds at low cost.

The objective of this study was to evaluate whether the evaluation of a transgene effect using DH lines, F_2:3_ lines, and F_3_ bulks would produce the same results across multiple environments and genetic backgrounds. A secondary objective was to evaluate whether the effect of the target transgene interacted with genetic background and environment.

## MATERIALS AND METHODS

### Germplasm

Through transgenesis, a native maize gene, the Zmm28 transcription factor (Wu et al. 2019), was overexpressed. A single event of the Zmm28 transgene was subsequently backcrossed into Corteva proprietary Stiff Stalk (SS) and Non-Stiff Stalk (NSS) heterotic group inbred lines. Using the backcross derived lines, two SS and two NSS families were generated as follows. Two families were created within each heterotic group by first crossing a transgenic inbred to two different, non-transgenic, inbred lines, resulting in hemizygous F_1_ seed. From the hemizygous F_1_ seeds, doubled haploid lines, F_2:3_ lines, and F_3_ line bulks were generated (Table 1). The DH lines were induced directly from the F_1_ parents using standard procedures (Röber et al., 2005), after which DH lines positive and negative for the transgene were identified. To create the F_2:3_ lines, several F_1_ plants of a given cross were self-pollinated and bulk harvested. The following growing season, F_2_ plants homozygous for the transgene or for the absence of the transgene were self-pollinated and individual ears were harvested, generating F_2:3_ lines with and without the transgene. The F_3_ bulks were generated by compositing equal quantities of seed from the individual F_2:3_ lines from each family with and without the transgene.

**Table 1.** The number of transgenic and non-transgenic doubled haploid (DH) lines and F_2:3_ lines and the number of replicate F_3_ bulk plots evaluated from two families in each of two heterotic groups

| Family | Heterotic Group | Transgenic | Non-transgenic | Transgenic | Non-transgenic | Transgenic | Non-transgenic |
| --- | --- | --- | --- | --- | --- | --- | --- |
|  |  | Doubled Haploid Lines |  | F<sub>2:3</sub> Lines |  | F<sub>3</sub> Bulk |  |
| DB30 | NSS | 20 | 18 | 11 | 17 | 13 | 14 |
| RO94 | NSS | 20 | 15 | 14 | 20 | 14 | 15 |
| WM45 | SS | 14 | 13 | 19 | 16 | 18 | 17 |
| JH61 | SS | 19 | 13 | 16 | 11 | 19 | 18 |

For trait evaluation, all lines were topcrossed to an inbred line tester from the opposite heterotic group. For each DH line, twenty-five plants were topcrossed and their seed bulked for testing. For each F_2:3_ line, seed was planted in several rows, topcrossed, and harvested in bulk. Twenty-five seeds from each F_3_ bulk were also planted and topcrossed, and each plant was harvested individually. A balanced bulk of the seed from these plants was used for hybrid testing of the F_3_ bulk. Approximately the same number of transgenic and non-transgenic lines were tested for each family for the DH and F_2:3_ methods; for the F_3_ bulks, a similar number of replicate plots was planted (Table 1).

### Field Trial Design

Field trials were conducted in 2017 at Woodland and Madison, CA. The Madison trial (MA26W) and one trial at Woodland (WO53W) were fully irrigated. A second trial at Woodland (WO71G) had water limitations imposed during the grain filling period, resulting in a mild stress and a fifteen percent yield reduction compared to the average of the two other locations. Other than water stress, all environments were managed with standard agronomic practices. The SS and NSS families were grown in separate experiments, with five replications per experiment for each environment. All entries (Table 1) were planted as two-row plots 4.9 m long with rows separated by 0.75 m. All plots were planted at 94,000 plants per hectare. The entire plot area was bordered with maize to minimize edge effects. All experiments were designed using a randomized complete block design, with replication as the blocking factor. All transgenic and non-transgenic lines, regardless of line creation method, were completely randomized within each replication.

### Data Collection

Grain yield, grain test weight, grain moisture, plant height, and ear height phenotypes were measured in all environments. Ear height was measured to the ear node and as the average of four randomly chosen plants within each plot, while excluding the three edge plants at each end of the plot. Plant height was measured to the collar of the flag leaf and was taken as the average of the same four randomly selected plants as was used for ear height. Ear height and plant height were only measured in two reps at each location. Grain yield, moisture, and test weight were measured on all plots using a two-plot combine.

### Statistical Analysis

All analyses were done using the ASREML software (Gilmour et al. 2015). All traits were analyzed using a mixed linear model with the transgene presence/absence (T), family (F), location (L), and line generation method (M) considered as fixed effects, while the replications within locations and individual line effects within family were considered random effects. The T × M, T × F, T × L, M × F, M × L, F × L, T × M × F, T × M × L, L × M × F, T × F × L, and T × F × L × M interaction terms were also included in the model. Three- and four-way interaction terms were small and non-significant, so the mixed-model was re-run without including them. The reduced models were evaluated to assess interactions for the remaining terms. Auto-regressive spatial adjustments were done for rows and columns within each location (Gilmour et al. 1997). Significance of fixed effects were assessed at the five percent probability level.

Best linear unbiased predictions (BLUPs) were generated for each entry across locations. The transgene effect for each trait for a given family was estimated as the difference in the trait value between transgenic and non-transgenic entries. In order to generate a variance of the predicted value, we used a line resampling process for the DH, F_2:3_, and F_3_ hybrids as follows. First, we sampled one line pair, computed the difference between transgenic and non-transgenic lines as described above, and repeated the process 1000 times, after which we computed the variance for the difference in transgene value for one line pair. We then repeated the process by randomly sampling two line pairs, computing the difference, repeating the process 1000 times, and computing the variance. We continued this process for 3 through 11 pairs. For the F_3_ hybrids, the difference was estimated by treating different plots within replications as separate entries, each representing different samples from the same F_3_ bulk. Because only 11 hybrids were evaluated for two families (Table 1), we used a maximum of 11 pairs to estimate the variance for the transgene effect for all families. The variances for different numbers of line pairs were then plotted for all families from each of the methods tested (Fig. 1).

**Figure 1.**
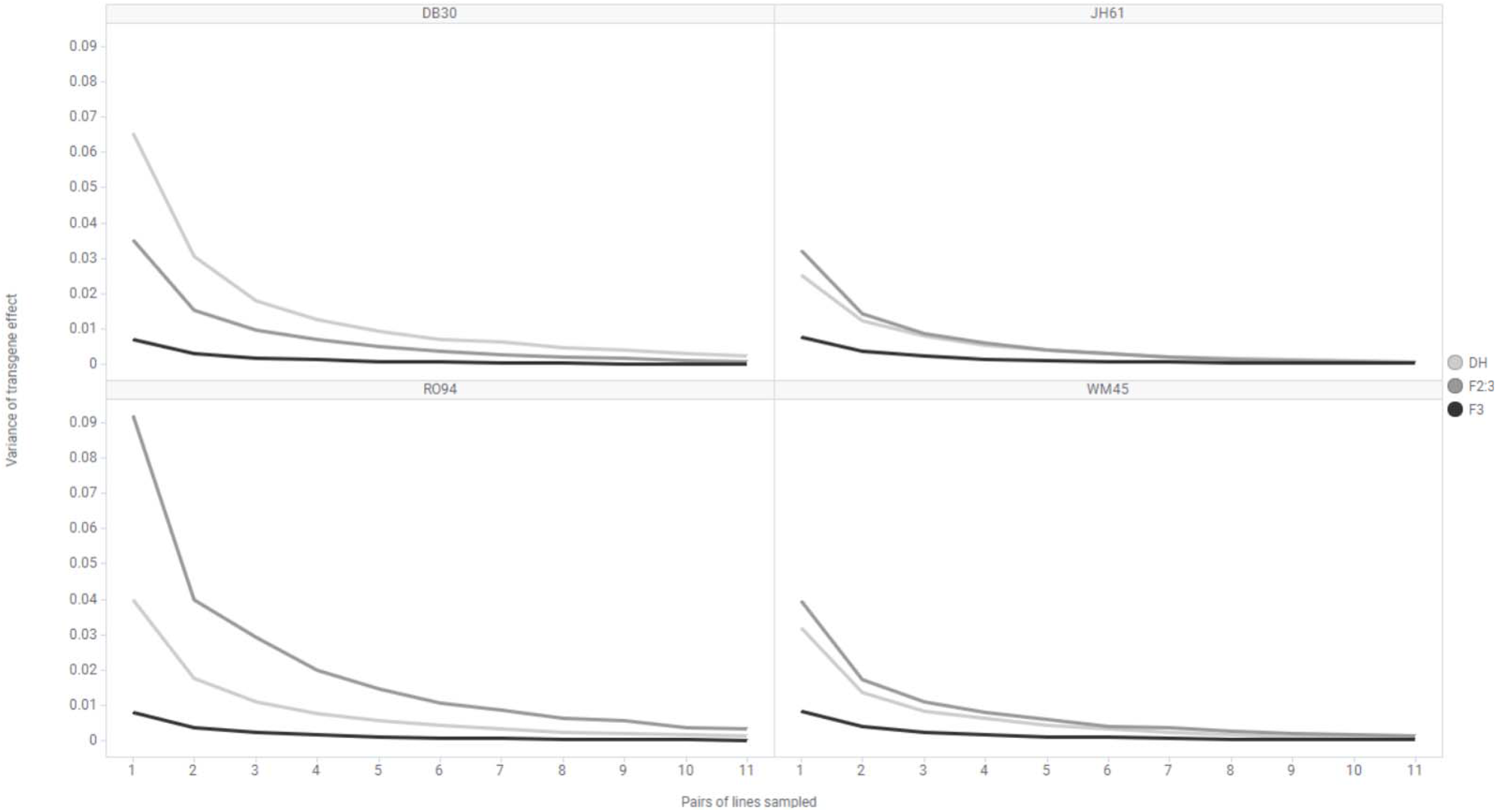
Variance of the estimated transgene effect for yield of testcross hybrids of two stiff stalk (SS) and two non-stiff stalk (NSS) families, based on randomly sampling 1 to 11 transgenic and non-transgenic pairs of DH or F_2:3_ hybrids or replicate plots of F_3_ families over 1000 sampling iterations.

## RESULTS

The transgene effect was significant for all traits except for plant height in the NSS families (Table 2). The line generation methods differed for yield in both heterotic groups and for moisture and test weight for the SS heterotic group. The family effects were mostly significant, except that the SS families showed no difference for yield and plant height. Locations differed for all traits for both heterotic groups.

**Table 2.**
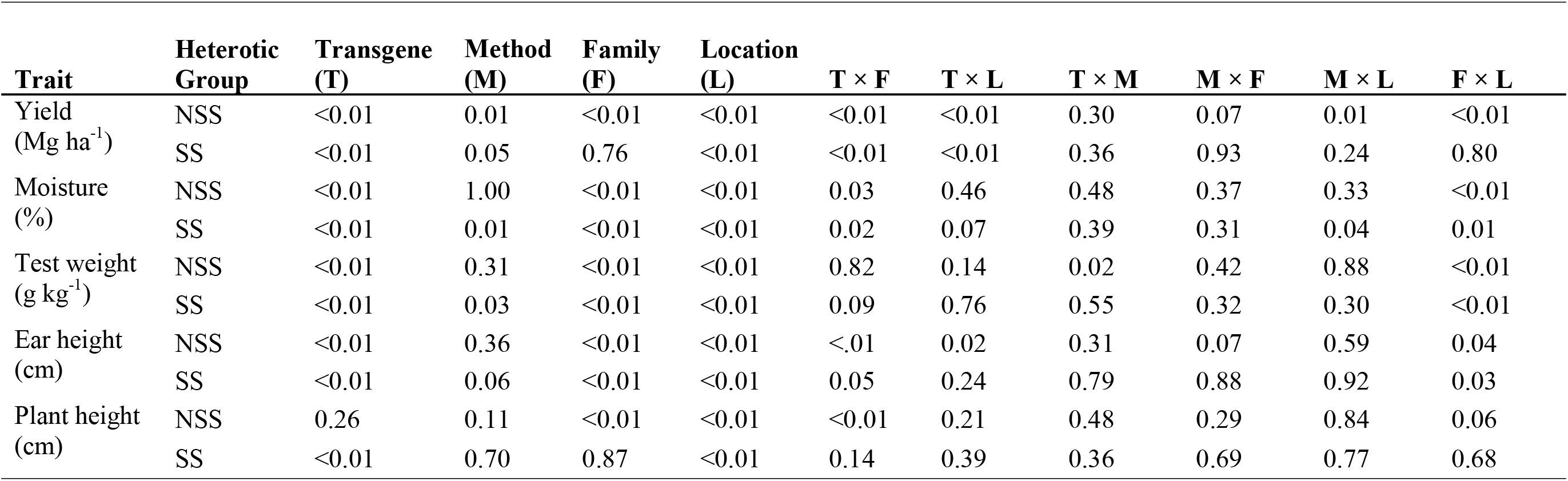
Probability values of F-tests of fixed effects of transgene (T), line generation method (M), family (F), and locations (L) and their two-way interactions for five maize traits measured on testcross hybrids of lines from two heterotic groups in three environments.

| **Trait** | **Heterotic Group** | **Transgene (T)** | **Method (M)** | **Family (F)** | **Location (L)** | **T × F** | **T × L** | **T × M** | **M × F** | **M × L** | **F × L** |
| --- | --- | --- | --- | --- | --- | --- | --- | --- | --- | --- | --- |
| Yield (Mg ha<sup>-1</sup>) | NSS | <0.01 | 0.01 | <0.01 | <0.01 | <0.01 | <0.01 | 0.30 | 0.07 | 0.01 | <0.01 |
|  | SS | <0.01 | 0.05 | 0.76 | <0.01 | <0.01 | <0.01 | 0.36 | 0.93 | 0.24 | 0.80 |
| Moisture (%) | NSS | <0.01 | 1.00 | <0.01 | <0.01 | 0.03 | 0.46 | 0.48 | 0.37 | 0.33 | <0.01 |
|  | SS | <0.01 | 0.01 | <0.01 | <0.01 | 0.02 | 0.07 | 0.39 | 0.31 | 0.04 | 0.01 |
| Test weight (g kg<sup>-1</sup>) | NSS | <0.01 | 0.31 | <0.01 | <0.01 | 0.82 | 0.14 | 0.02 | 0.42 | 0.88 | <0.01 |
|  | SS | <0.01 | 0.03 | <0.01 | <0.01 | 0.09 | 0.76 | 0.55 | 0.32 | 0.30 | <0.01 |
| Ear height (cm) | NSS | <0.01 | 0.36 | <0.01 | <0.01 | <.01 | 0.02 | 0.31 | 0.07 | 0.59 | 0.04 |
|  | SS | <0.01 | 0.06 | <0.01 | <0.01 | 0.05 | 0.24 | 0.79 | 0.88 | 0.92 | 0.03 |
| Plant height (cm) | NSS | 0.26 | 0.11 | <0.01 | <0.01 | <0.01 | 0.21 | 0.48 | 0.29 | 0.84 | 0.06 |
|  | SS | <0.01 | 0.70 | 0.87 | <0.01 | 0.14 | 0.39 | 0.36 | 0.69 | 0.77 | 0.68 |

A significant T × F interaction was identified for most traits, except for test weight and plant height within the SS heterotic group. A significant T × L effect was present in both heterotic groups for yield, but otherwise, only for ear height in the SS heterotic group. The only significant T × M interaction was for test weight for the NSS families (Table 2), where the F_3_ derived transgenic hybrids were 4 kg m^−3^ lower than non-transgenic families, but no difference was detected between transgenic and non-transgenic hybrids originating from DH lines. A M × F interaction was not detected for any trait. The M × L interaction was significant for yield in the NSS heterotic group and for moisture in the SS heterotic group but absent otherwise (Table 2). The 3 and 4-way interactions were non-significant or small and are not discussed further.

Overall, the transgene had a positive effect on yield, with an average increase of 0.3 Mg ha^−1^ across both the NSS and SS families (data not shown). Moisture at harvest was higher for transgenic hybrids for both NSS, by 0.1 percentage points, and SS, by 0.3 percentage points, heterotic groups. Test weight in non-transgenic hybrids was consistently higher by 2 kg m^−3^ in NSS families and 5 kg m^−3^ in SS families. Transgenic hybrids had higher ear heights, averaging 3 cm taller for NSS and 6 cm taller for SS families compared to their non-transgenic comparators. There were no differences between transgenic and non-transgenic hybrids of NSS families for plant height. However, transgenic hybrids of SS families were on average 4 cm taller than non-transgenic hybrids (data not shown).

The effect of the transgene varied among families for most traits (Table 3). The transgene increased yield in both the DB30, by 0.5 Mg ha^−1^, and WM45, by 0.7 Mg ha^−1^, families but no effect in families RO94 or JH61 (Table 3). For moisture, the transgenic hybrids had higher moisture at harvest in both SS families but the increase in family WM45 was 0.2 percentage point more than for family JH61. For the NSS families, there was no difference in grain moisture at harvest between transgenic and non-transgenic hybrids in family DB30. However, the transgenic hybrids in family RO94 had 0.2 percentage points more moisture than the non-transgenic hybrids. Transgenic hybrids in family RO94 were 3 cm shorter, but ears were 2 cm higher than non-transgenic hybrids. In contrast, in family DB30, transgenic hybrids were 2 cm taller than non-transgenic hybrids and increased ear height by 2 cm more than in the RO94 family. In the SS families, transgenic hybrids increased ear height by 5 cm for the JH61 family and 6 cm for the WM45 family.

**Table 3.** Best linear unbiased predictions for four traits measured on testcross hybrids of transgenic and non-transgenic lines from the Non-Stiff Stalk and Stiff Stalk maize heterotic groups across three environments

| Trait | Transgenic Hybrid | Non-Transgenic Hybrid | Difference | Transgenic Hybrid | Non-Transgenic Hybrid | Difference |
| --- | --- | --- | --- | --- | --- | --- |
| Non-Stiff Stalk Families |  |  |  |  |  |  |
|  | DB30 |  |  | RO94 |  |  |
| Yield (Mg ha<sup>-1</sup>) | 14.4 | 13.9 | 0.5* | 13.5 | 13.4 | 0.1 |
| Moisture (%) | 16.2 | 16.3 | 0.1 | 15.6 | 15.4 | 0.2* |
| Plant Height (cm) | 289 | 287 | 2* | 264 | 267 | -3* |
| Ear Height (cm) | 147 | 143 | 4* | 134 | 132 | 2* |
| Stiff Stalk Families |  |  |  |  |  |  |
|  | JH61 |  |  | WM45 |  |  |
| Yield (Mg ha<sup>-1</sup>) | 14.8 | 14.8 | 0.0 | 15.1 | 14.4 | 0.7* |
| Moisture (%) | 16.7 | 16.5 | 0.2* | 17.2 | 16.8 | 0.4* |
| Plant Height (cm) | 317 | 314 | 3* | 317 | 313 | 4* |
| Ear Height (cm) | 166 | 161 | 5* | 164 | 158 | 6* |

\*Difference between the transgenic and non-transgenic hybrids is significant at the .05 probability level.

Any given pair of DH or F_2:3_ lines with and without the transgene represent one possible comparison for the population of line pairs. Any given pair of lines is developed arbitrarily; they are not near-isogenic comparisons. Therefore, we estimated the variance among line pairs in their transgene effects to determine the number of line pairs that would need to be evaluated to get an accurate estimate of the transgene effect in the population. We sampled from 1 to 11 hybrid pairs from the DH, F_2:3_ and F_3_ generations. Across all 4 families, the F_3_ hybrids consistently required fewer pairs to achieve a stable estimate of a transgene effect, with even a single pair showing minimal variance (Figure 1), as might be expected since every “pair” is a replicate of the same set of bulked lines. The DH and F_2:3_ generations consistently required more pairs to achieve the same low variance as a single pair of the F_3_ generation. For the NSS families it was necessary to sample 5 times more hybrid pairs for F_2:3_ and DH generations to achieve a stable estimate, while 3 times the number of hybrid pairs was required for SS families to achieve the same low variance as F_3_ hybrid pairs.

## DISCUSSION

In this experiment, we found that the transgene effect was not influenced by the method of line evaluation. We do not believe this is due to a lack of power in our analysis, because differences among hybrids were detectable at the transgene main effect level across locations if they were equal to or larger than 0.1 Mg ha^−1^ for yield, 1.3 kg m^−3^ for test weight, and 0.1 percentage points for moisture. Differences could be detected at 1 cm for plant height and ear height, which were both measured only on two of the five available replications. Thus, we feel confident that we could identify the effect of a transgene by any of the three methods we evaluated here.

The Zmm28 transgene’s main effect was conditional on other factors, including location and genetic background (i.e., family). These results reinforce previous studies that demonstrated the significant impact of both the genetic background and environment on the estimated effect of a transgene for quantitative traits (Linares, et al., 2022; Linares, 2021). Consequently, because breeding programs need to sample both environments and germplasm in order to assess the value of a transgene for a quantitative trait, our objective was to determine the best type of family to use to conduct this assessment as rapidly and at as low cost as feasible. We hypothesized that different line generation methods would provide a similar ability to estimate a transgene’s value, and therefore, methods that have lower cost could be prioritized. We found no evidence of a consistent interaction of line generation method with family and only one instance of a transgene by method interaction. Thus, testing the Zmm28 event in topcrosses of DH lines, F_2:3_ families, or F_3_ family bulks all provided a similar assessment of the transgene value. However, to assess the transgene effect on the population, F_3_ family bulks required the fewest resources, because all within family variation was captured in a single plot such that only one transgenic and one non-transgenic plot was needed for each population (which could be replicated if desired). However, for the other methods, multiple positive and negative lines were required to test the transgene effect.

The use of DH lines in most commercial maize breeding programs makes them an initially obvious choice in which to assess the value of a transgene intended to affect a quantitative trait. They have many advantages, including being homozygous, stable lines that can be repeatedly tested across years and directly incorporated into breeding programs. Use of DH lines can also permit the evaluation of the transgene’s effect on inbred line seed production. However, DH lines are costly and take time to be developed and evaluation of a transgene’s effect will require testing multiple transgenic and non-transgenic lines. To save time, F_2:3_ lines could be used as a proxy for DH lines, enabling estimations of within family variance without the extra time and expense necessary for DH line creation. Evaluation of transgene positive and negative lines would still require multiple plots to determine the overall transgene effect in the population.

Many transgenes designed to impact qualitative traits are directed towards single protein targets and tend to be highly penetrant and have consistent effect across the germplasm for the targeted trait. In contrast, as has been observed for QTL (e.g., Boer et al. 2007), transgenes for quantitative traits are more prone to transgene by family by environment interactions necessitating broader testing to assess their value (Guo et al. 2014, Simmons et al. 2021). Therefore, we argue that at the early stages of testing, using the older method to create F_2:3_ lines and then bulking them to form F_3_ positive and negative bulks offers the fastest way to determine whether a transgene has value for a given quantitative trait across many diverse populations within the breadth of a breeding program’s germplasm.

While transgene effects were largely consistent across the three methods, we observed an interaction between the transgene and the line generation method (T × M), for test weight in the NSS heterotic group. The DH method showed no transgene effect, compared to a 4 kg m^−3^ effect in the F_3_ hybrids. One possible explanation for this result could be the reduced number of recombination events sampled in the F_1_ derived DH lines used in this study, which would result in large linkage blocks (Sleper and Bernardo, 2016). Up to 25 percent of the phenotypic variation for test weight in maize could be attributed to five QTL (Ding et al., 2011) in a single population, which further reinforces the important influence of recombination events if using DH lines for evaluation of transgenes. This effect was not detected for the SS heterotic group families.

The F_3_ bulking approach could have some limitations. Strong within plot competition among individual plants could act to obscure a positive transgene effect. For example, in the case of a phenotype such as reduced stature that increases yield, shorter plants within a plot, although having increased yield potential, could be at a competitive disadvantage for intercepting incoming solar radiation in a bulked planting. Ultimately, the influences of environments and genetic backgrounds on the effect of a transgene (or a QTL) necessitate a diverse testing plan. Here, we demonstrated that three different line generation methods generally gave equivalent results when measuring the effect of a transgene across a sample of breeding germplasm. If the breeding goal is to assess the transgene value for the breeding program at large, rather than for a specific genetic background, then the use of F_3_ bulks appears to be the most useful method.

## Abbreviations

QTL: Quantitative Trait Locus/Loci

## ORCID

J.F. Linares: 0000-0002-0083-0974); N.D. Coles: 0000-0002-2008-3283; J.E. Habben: 0000-0001-6609-153X); C. Messina: 0000-0002-5501-9281; M. Cooper: 0000-0002-9418-3359; R. Carrasco: 0000-0001-5702-1396; Javier Carter: 0000-0003-3287-9956; and E.C. Brummer: 0000-0002-3621-4516.

